# Spin-Dependent Extracellular Respiration

**DOI:** 10.64898/2026.05.29.728850

**Authors:** Nir Sukenik, Cole C. Harris, Sukrampal Yadav, Marko Chavez, Christina M. Niman, Lech Tomasz Baczewski, Mohamed Y. El-Naggar

**Author notes:** These authors contributed equally to this work (Nir Sukenik, Cole C. Harris, Sukrampal Yadav, Marko Chavez).

## Abstract

Biological energy conversion relies on highly efficient electron transfer. The chirality induced spin selectivity (CISS) effect, which couples electron spin to momentum in chiral molecules, is hypothesized to promote this efficiency. While observed in isolated biomolecules, the physiological relevance of CISS during active cellular metabolism remains unknown. Here, we demonstrate that CISS influences extracellular electron transfer in living *Geobacter sulfurreducens* biofilms. Cultivation on ferromagnetic electrodes yields a significant asymmetry in respiratory current between opposite substrate spin states. Furthermore, *in situ* magnetization reversal induces reversible changes in respiratory flux. These results provide the first *in vivo* demonstration that spin selectivity directly impacts respiration. By revealing a quantum feature of extracellular respiration, our findings offer a strategy to exploit the spin degree of freedom in bioelectronics.

## Introduction

Electron transfer (ET) drives the energy transduction pathways of living cells, including respiration and photosynthesis. Within respiratory chains, electrons derived from metabolic substrates are channeled through a sequence of protein-embedded redox cofactors, ultimately reducing a terminal electron acceptor to power cellular metabolism. Remarkably, these sequential ET processes exhibit high fidelity, facilitating efficient and directional charge transport along specific biomolecular pathways. Despite spanning a nanometer-to-micrometer distance through organic frameworks typically regarded as electrical insulators, this specificity prevents short-circuit reactions and the off-path diffusion of high-energy redox equivalents. Such precision is a biological imperative; diverting electrons from productive pathways is thermodynamically wasteful, and risks generating destructive moieties (*1*–*4*). A defining structural characteristic of these biological electron transport chains is the chirality of the charge-mediating proteins, an asymmetry that propagates from individual homochiral L-amino acid monomers through higher-order secondary structures. While the origin of this biological homochirality may itself be rooted in spin-selective precursor reactions (*5, 6*), its universality across extant life raises a fundamental question: Does this structural asymmetry promote the extraordinary efficiency of biological electron transport?

A recent hypothesis links chirality and efficient ET through the chirality induced spin selectivity (CISS) effect (*7, 8*). CISS couples an electron’s spin to its linear momentum as it traverses a chiral electrostatic potential. Consequently, electron transmission becomes spin-dependent, favoring one spin orientation (e.g., parallel or anti-parallel to its momentum) dictated by molecular handedness. While the theoretical basis of the CISS effect is intensely debated, its prevalence in chiral systems and implications for spintronics, enantioselective chemistry, and biochemical processes are now widely demonstrated (*8*). In biological electron transport chains, the forward (productive) direction of ET is fundamentally defined by the thermodynamic driving force: the overall gradient of redox midpoint potentials spanning from electron donor to terminal acceptor. Navigating this cascade often requires traversing local energetic barriers that can exacerbate reverse electron tunneling and charge recombination, requiring careful control of redox site spacings and electronic couplings (*3, 9, 10*). This is where the spin–momentum coupling of CISS can become physiologically relevant: it inherently suppresses backward transitions for the favored spin channel (*7, 8*). By rendering reverse ET steps spin-restricted, CISS ensures that the overall redox driving force translates to unidirectional, forward propagation. Within biological ET networks, CISS implies that homochiral protein assemblies systematically bias charge transport, enhancing forward transfer rates and overall efficiency.

Over the past decade, the CISS effect has been definitively established across a wide range of chiral systems, including isolated biomolecules such as DNA, peptides, and proteins (*8, 11*–*15*). However, these demonstrations primarily probe isolated molecules under non-physiological conditions, such as dehydrated monolayers subjected to high driving voltages. These settings remain far removed from the aqueous environments and physiological redox windows of living cells. Thus, a critical question remains unresolved: Does spin selectivity also functionally dictate electron transport and respiration *in vivo*? Resolving this demands an experimental design that combines direct, quantitative monitoring of cellular respiration with external control over the spin state of a terminal electron acceptor.

The extracellular respiration pathway of certain electroactive bacteria uniquely fulfills these biophysical requirements by exporting electrons beyond the cell membrane, offering a direct electrical readout of cellular respiration alongside an accessible interface for external spin-state manipulation (*16*). Instead of reducing soluble oxidants that diffuse inside the cells, these organisms gain energy for growth using an extracellular electron transfer (EET) pathway that connects the intracellular electron transport chain to the reduction of insoluble terminal electron acceptors, such as iron or manganese oxide minerals located extracellularly (*17*). The EET pathway of *Geobacter sulfurreducens*, one of the most studied model electroactive bacteria, consists of a network of multiheme cytochromes spanning the periplasm, outer-membrane, and extending into extracellular nanowires for electron transport from the inner membrane to the cell surface and beyond (*16, 18*–*22*). The CISS effect was recently demonstrated in purified cytochromes from another electroactive bacterium (*Shewanella oneidensis*) (*13, 14*). While these measurements were limited to non-physiological conditions, they motivate the question of whether the CISS effect is operative during EET *in vivo. G. sulfurreducens* can utilize solid-state electrodes as the terminal electron acceptor for respiration, resulting in high EET currents, cellular growth, and the formation of thick multilayered biofilms on the electrodes (*23*–*25*). Consequently, the overall respiration rate of the biofilm can be directly quantified, in real time, as an electrochemical current to a working electrode. During active metabolism, biofilm voltammetry exhibits a reversible catalytic wave that reaches a limiting current by -0.34 V vs. Ag/AgCl (*23*). By employing ferromagnetic (FM) working electrodes with out-of-plane magnetic anisotropy, where the spin states are well defined by a strong exchange coupling, the spin orientation of the electron acceptor can be externally controlled by reversing the magnetization direction by using an external magnet. This architecture enables direct, *in vivo* interrogation of spin-dependent respiration (Fig. 1).

**Fig. 1:**
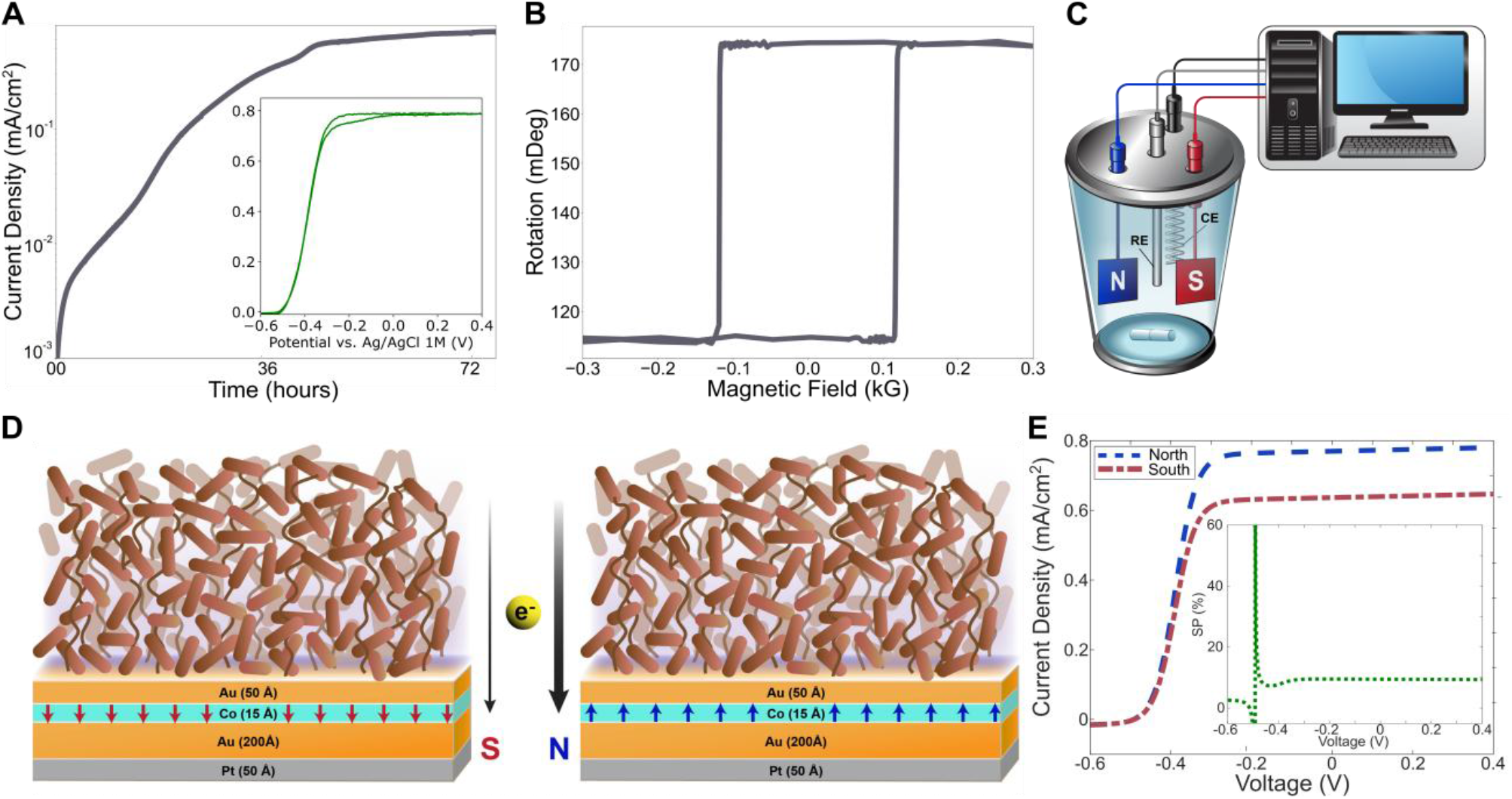
Magneto-bioelectrochemical approach for probing spin-dependent extracellular respiration. **(A)** Representative semi-logarithmic chronoamperometry (CA) profile of *Geobacter sulfurreducens* biofilm growth on a working electrode. (Inset) Representative cyclic voltammogram (CV) exhibiting catalytic current generation after the biofilm reached maximal CA current density. **(B)** Polar magneto-optic Kerr effect (MOKE) hysteresis loop of the ferromagnetic (FM) electrodes, demonstrating strict out-of-plane magnetic anisotropy with a 117 G coercive field. **(C)** Schematic of the electrochemical reactor containing a reference electrode (RE), counter electrode (CE), and two oppositely magnetized FM working electrodes (N and S). **(D)** Conceptual illustration of spin-selective extracellular electron transfer (EET) to FM electrodes. The chiral biological network selectively filters electrons according to the chirality induced spin selectivity effect, favoring higher biofilm EET currents for the electrode with the preferred spin state. **(E)** Theoretical Nernst-Monod projection of CVs for N and S electrodes exhibiting distinct kinetic parameters. (Inset) Expected potential-dependent spin polarization, *SP(E)*, calculated from eq. 1 using currents derived from the theoretical Nernst-Monod CVs.

Here, we demonstrate that the spin orientation of the terminal electron acceptor modulates the respiration of living *G. sulfurreducens* biofilms. Biofilms cultivated on FM electrodes pre-magnetized in opposite directions exhibit an asymmetry in current output. Chronoamperometry (CA) and cyclic voltammetry (CV) measured during active cellular metabolism, where acetate oxidation drives a sustained electron flux to the electrode, reveal elevated EET current for the biologically preferred magnetization orientation, yielding a stable spin polarization (SP) of 10.5 ± 1.6 %. Furthermore, *in situ* magnetization reversal of pre-established biofilms induces rapid, reversible modulation of the respiratory current. These results provide *in situ* evidence that the electron spin state gates extracellular respiration in a manner consistent with the CISS effect, establishing a physiological role for spin-dependent processes in living systems.

## Results

### Experimental Design: A magneto-electrochemical approach to investigate spin-dependent extracellular electron transfer

To establish whether a spin-dependent phenomenon actively impacts *in vivo* extracellular respiration, we developed a magneto-bioelectrochemical platform to continuously monitor the EET current of *G. sulfurreducens* biofilms (Fig. 1). Following inoculation, CA reveals a rising anodic current density tracking cellular acetate oxidation and biofilm growth, which stabilizes at a maximum mass-transport-limited current (Fig. 1A) (*23, 24, 26*). Subsequent CV exhibits a catalytic wave at the formal potential of the EET cytochromes (Fig. 1A inset). To test the role of spin, we grew epitaxial Au/Co/Au ferromagnetic (FM) electrodes with a well-defined out-of-plane magnetic anisotropy via molecular beam epitaxy. Polar magneto-optic Kerr effect (MOKE) measurements reveal a sharp, rectangular hysteresis loop with a 117 G coercive field (Fig. 1B), indicating unidirectional perpendicular alignment of the cobalt spins. A biocompatible 5 nm gold capping layer prevents oxidation of the underlying 1.5 nm cobalt layer. Using an external magnet, we pre-magnetized the electrodes into spin-up (N) or spin-down (S) magnetized states. Simultaneously cultivating biofilms on oppositely magnetized electrodes within the same reactor (Fig. 1C) isolates the impact of electron spin and eliminates culture-to-culture variability.

If biofilm EET is spin-selective, we reasoned that the N vs. S comparison would yield an asymmetry due to CISS: enhanced current density and catalytic activity for the biologically preferred magnetization state (Fig. 1D). We project this asymmetry using the established Nernst-Monod model of microbial biofilm voltammetry (*26*). An observed spin-dependent effect will manifest as distinct kinetic parameters for the two magnetization states – specifically, a difference in the maximum limiting current density and the apparent half-saturation potential (Fig. 1E). In the context of CISS studies, the potential dependent spin polarization (*SP(E)*) is typically defined as the normalized difference in current density for the two magnetization states at each potential, as shown in eq. 1 (*8*).

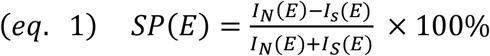

Fig. 1E inset depicts a theoretical expected *SP(E)* response calculated for two Nernst-Monod curves with distinct kinetic parameters. This approach establishes a clear profile for the expected live cell signal.

### Spin-Dependent Measurements of Biofilm Catalytic Activity

Strikingly, CV scans across independent reactors reveal consistently higher catalytic current densities for biofilms respiring N electrodes relative to their S counterparts (Fig. 2, A to C). Table S1 contains growth conditions and CV measurement parameters for all replicate reactors. The potential-dependent *SP(E)*, calculated from the forward scans (Fig. 2D) matches the expected kinetic asymmetry profile (Fig. 1E). Above potentials required to support EET (-0.34 V vs. Ag/AgCl), *SP(E)* stabilizes to a steady plateau with a highly reproducible magnitude (10.5 ± 1.6 %) across independent reactors despite varying baseline current densities (Fig. 2, A to C). Furthermore, fitting the CV data (fig. S1 and table S2) reveals a corresponding difference in the Nernst-Monod kinetic parameters between the two oppositely magnetized electrodes. In contrast, control experiments utilizing identically magnetized (N-N) electrode pairs yielded similar CA current and hence a negligible SP (Fig. 2D and fig. S2B), confirming that the observed N-S asymmetry arises from the opposite magnetization states of the electrodes.

**Fig. 2:**
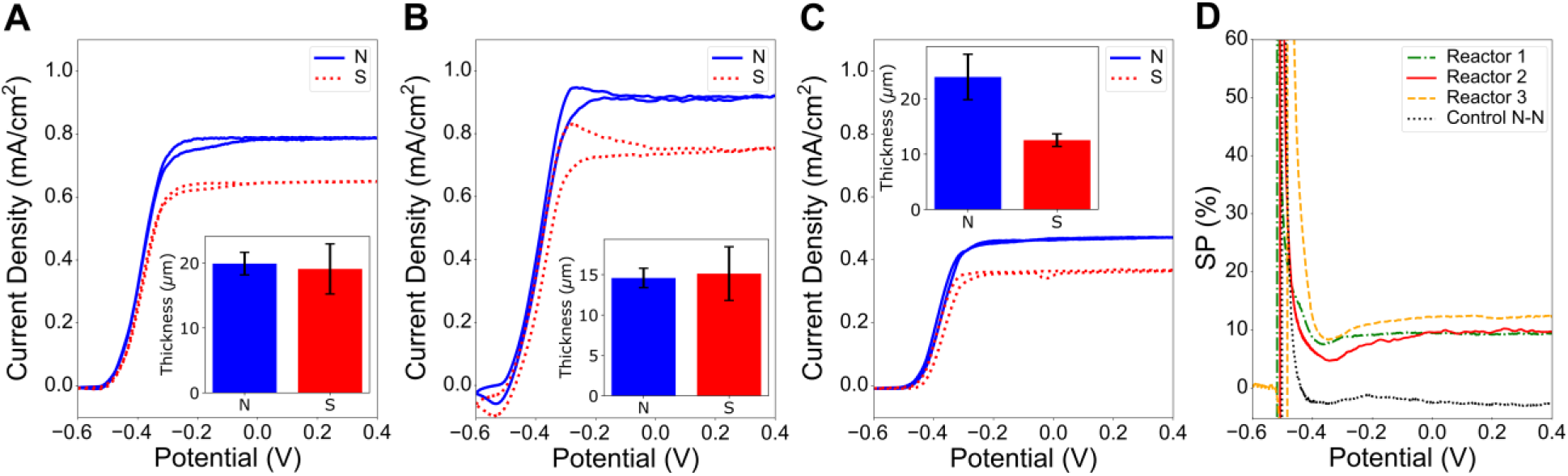
Spin-dependent catalytic activity of *G. sulfurreducens* biofilms. **(A to C)** CVs comparing the catalytic current densities of biofilms cultivated simultaneously on N-magnetized (solid blue) and S-magnetized (dotted red) ferromagnetic electrodes across three independent reactors (Reactors 1-3 in A to C, respectively). The N electrodes consistently support enhanced catalytic current. (Insets) Average biofilm thicknesses measured via confocal microscopy at the end of each experiment, showing comparable biomass for saturated biofilms (A and B) and higher biomass for the N electrode when terminated during exponential growth (C). Error bars represent standard deviation. **(D)** Potential-dependent *SP(E)*, calculated from the forward CV scans of the three reactors compared to a control reactor containing two identically magnetized (N-N) electrodes. The N-S pairs converge to a stable 10.5 ± 1.6% physiological *SP(E)* plateau at applied potentials supporting EET, whereas the N-N control yields a negligible response.

To determine if the enhanced catalytic activity on N electrodes and the resulting SP signal stem from an increase in total biofilm thickness or an intrinsic enhancement of overall EET activity per unit biomass, we measured biofilm thickness for N vs. S electrodes using confocal laser scanning microscopy (table S3). Mature N and S biofilms reaching maximum plateau currents exhibited comparable thicknesses (Fig. 2, A and B insets). In this saturated regime, growth and activity become constrained by bulk mass transport limitations (e.g., nutrient diffusion) rather than electron transfer kinetics. However, when CA was terminated earlier during exponential current rise (a regime still governed by EET kinetics; fig. S2A), the favored N electrode accumulated biomass faster to yield a thicker intermediate biofilm (Fig. 2C inset). Consequently, the N magnetization supports a higher intrinsic EET activity that persists even after both biofilms reach mass-transport-limited mature thickness. After biofilm growth and turnover CV measurements, the reactors’ media were exchanged with fresh acetate-free media. When CV measurements were performed under these non-turnover conditions, no significant difference was observed in the biofilm redox peaks between N and S electrodes in all reactors (fig. S3). The spin-dependent asymmetry therefore emerges specifically during active respiration, coupling the oxidation of acetate and electron transport through the biofilm to the electrode.

### Tracking the Spin-Dependence of Extracellular Respiration During Biofilm Growth

Because current production by electroactive biofilms tracks metabolic activity, comparing the temporal current density, *I(t)*, between paired N and S electrodes across multiple reactors (Fig. 3, A to C) reveals the temporal evolution of spin polarization, *SP(t)*, during cellular growth, as follows:

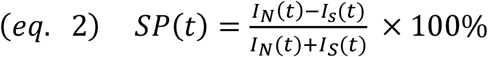

**Fig. 3:**
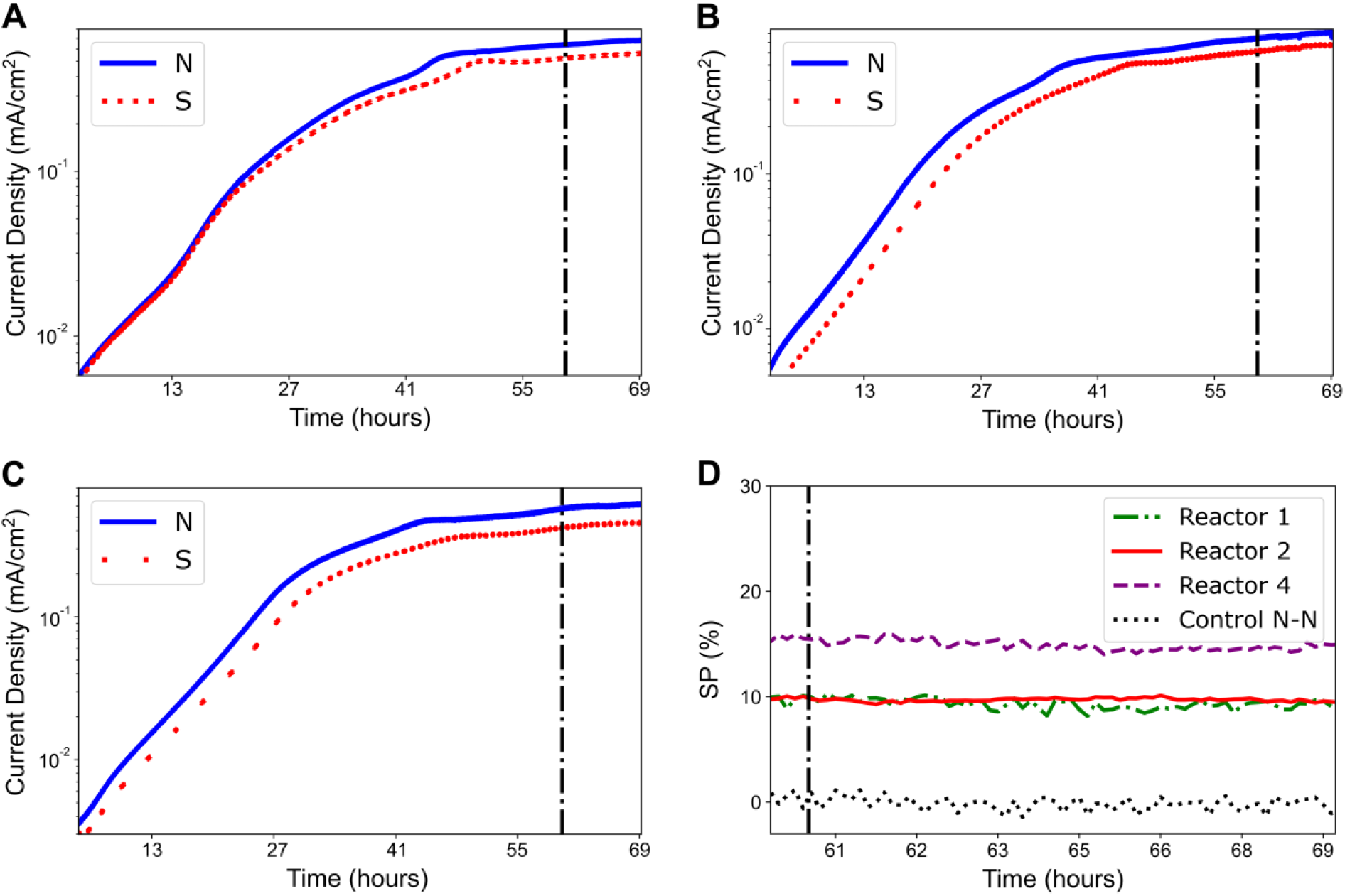
Temporal evolution of spin-polraized extracellular electron transfer during active biofilm growth. **(A to C)** Semi-logarithmic CA current profiles monitoring the growth of *G. sulfurreducens* biofilms on N-magnetized (solid blue) and S-magnetized (dotted red) electrodes in three independent reactors (Reactors 1, 2, 4 in A to C respectively). Vertical black dot-dashed lines indicate the onset of maximum current density from mature biofilms. A clear kinetic preference for the N-magnetized electrode emerges early during exponential growth and persists through biofilm maturation. **(D)** Time-dependent spin polarization, *SP(t)*, calculated for the mature biofilm stage for the three reactors and an identically magnetized (N-N) control reactor. The *SP(t)* signal maintains a stable plateau over extended multi-hour timescales, consistent with the steady-state *SP(E)* measurements, while the N-N control remains at baseline.

The semi-logarithmic *I(t)* profiles exhibit exponential growth followed by a plateau; these features mirror standard bacterial growth, and can be modeled with a logistic growth function (eq. S3, fig. S4 B to E, and table S4) (*27*–*29*). All measurements reveal a consistently higher respiratory current density for the N electrodes compared to their S counterparts, and this spin preference manifests well before biofilm activity reaches its stationary plateau (Fig. 3).

Early-time *SP(t)* signals exhibit complex dynamics (fig. S4A), reflecting stochastic initial cell attachment to the electrode. However, during late exponential growth, *SP(t)* settles to a constant long-term plateau (Fig. 3D) of 12.7 ± 3.7 % matching the 10.5 ± 1.6% *SP(E)* magnitude observed in mature biofilm voltammetry (Fig. 2). The spin preference and *SP(t)* trend are conserved across different applied growth potentials (Fig. 3 and Table S1). This result corroborates the flat profile of the *SP(E)* signal observed at all potentials higher than the formal potential of *G. sulfurreducens* EET (Fig. 2D).

### *In Situ* Modulation of Respiratory Current

While comparing separate N and S electrodes revealed a clear physiological spin preference emerging over multi-day growth periods (Fig. 2 and 3), we also sought to establish whether the terminal electron acceptor’s spin state continuously governs the respiratory flux of the same biofilm in real time. We executed a dynamic *in situ* magnetization reversal protocol on an established biofilm using an external magnet (Fig. 4A). The external magnet is only introduced temporarily to reorient the magnetization and immediately removed until the next reversal; this ensures that observed current modulations arise strictly from the electrode’s spin state, eliminating artifacts from the external magnetic field itself.

**Fig. 4:**
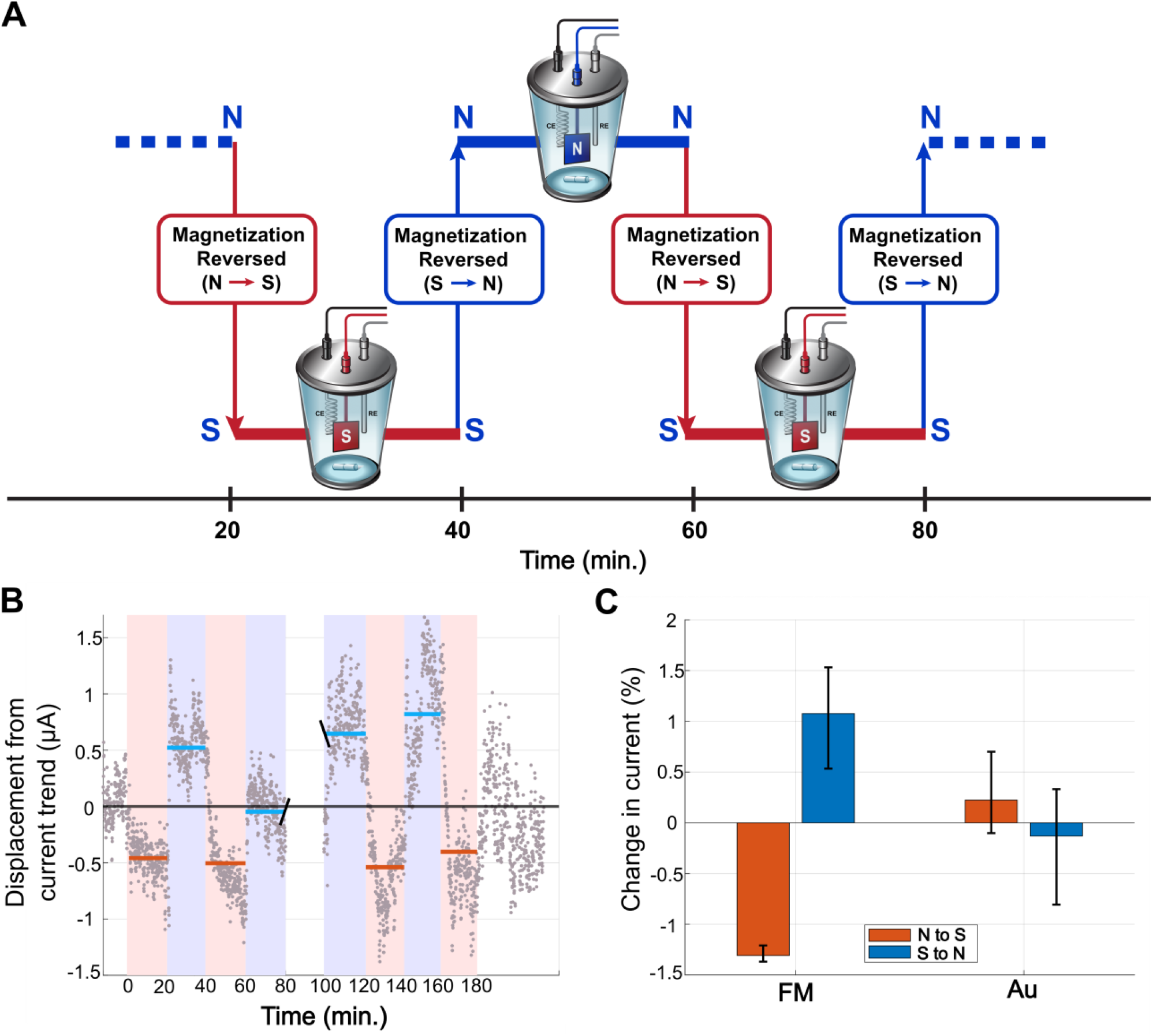
*In situ* modulation of respiratory flux by external magnetization reversal. **(A)** Schematic of the dynamic magnetization reversal protocol. A mature *G. sulfurreducens* biofilm is poised on the ferromagnetic (FM) electrode. An external (∼1000 G) magnet is momentarily introduced every 20 minutes to reverse the remanent magnetization between N and S states, then immediately removed to eliminate external magnetic field artifacts during active current monitoring. **(B)** Detrended CA current displaying rapid, reversible modulation in respiratory flux coupled to the electrode’s magnetization state. Shaded regions correspond to the N (blue) and S (red) magnetization intervals. Horizontal lines represent the mean current for each respective interval. **(C)** Average percentage change in respiratory current induced by N-to-S (red) and S-to-N (blue) magnetization reversals for the FM electrode and a non-magnetic gold (Au) control. Error bars indicate the full data spread, representing the dispersion in the detrended current response. The FM electrode exhibits a >1% modulation due to magnetization reversal, consistent with the preferred magnetization orientation from paired-electrode experiments, which are not observed in the Au electrode.

The detrended current reveals a rapid and reproducible modulation in EET flux following each magnetization reversal (Fig. 4B). While the magnitude of this current modulation (Fig. 4C) is smaller than the steady-state SP obtained from comparing N to S electrodes (Fig. 2D and 3D), it is significantly larger than the negligible baseline fluctuations of a biofilm cultivated on a gold only (non-magnetic) control electrode under similar conditions (fig. S5). Importantly, the sign of the response mirrors the paired-electrode experiments: switching to the N state instantaneously increases current production, whereas switching to the S state decreases it. This rapid, reversible switching provides *in vivo* evidence that the rate of extracellular respiration is continuously and dynamically gated by the spin state of the terminal electron acceptor.

## Discussion

The results presented here establish a preference for a specific electron spin orientation during extracellular respiration by living *G. sulfurreducens* biofilms. Importantly, our experimental design isolates a spin-dependent phenomenon unconfounded by external magnetic fields. While previous studies have reported altered currents in microbial electrochemical systems subjected to static magnetic fields (*30*–*32*), those responses were often ascribed to magnetohydrodynamic effects or Lorentz forces acting on ions (*33, 34*). By utilizing pre-magnetized thin-film FM electrodes with a well-defined out-of-plane magnetic anisotropy (Fig. 1), our experiments do not subject the biofilm to an external magnetic field during measurements. Because the macroscopically wide but ultra-thin perpendicularly magnetized cobalt layer acts as an infinite uniform magnetic sheet, it generates zero stray field above the electrode surface (*35*). This isolates a pure spin-dependent interfacial electron transfer effect, providing direct evidence for a preferred spin orientation associated with EET in a manner consistent with the CISS effect.

Remarkably, the physiological preference for the N electrode observed in these living biofilms is consistent with the sign of SP previously associated with electron transport measurements through dry, purified monolayers of multiheme cytochromes using magnetic conductive atomic force microscopy (*13*–*15*). The conservation of this preferred spin orientation, from isolated EET proteins in electrode-spanning molecular junctions to metabolically active biofilms, strongly indicates that the CISS effect is an intrinsic feature of electron transfer that bridges molecular transport and cellular-scale respiration.

Despite expected culture-to-culture variability in absolute current densities across independent reactors, the potential-dependent *SP(E)* (Fig. 2D), and the time-dependent *SP(t)* (Fig. 3D) profiles stabilize to a flat profile as the biofilms mature. Interestingly, however, the early stages of biofilm development following inoculation reveal more complex *SP(t)* dynamics (fig. S4A). Modeling the temporal current profiles with a logistic growth function (fig. S4, B to E) indicates that these dynamics are driven by the inherent stochasticity of initial cell attachment. Furthermore, evaluating the time derivative of current production (fig. S6) reveals consistently steeper rises for the N electrodes, reflecting a faster accumulation of electroactive biomass relative to their S counterparts. Ultimately, despite early temporal fluctuations, the spin preference dominates as the biofilm develops, driving all replicates to a *SP(t)* plateau of 12.7 ± 3.7 % when maximal current density is reached.

Notably, the measured SP signal likely underestimates the intrinsic biological spin preference because the finite thickness of gold between the chiral system and underlying ferromagnetic layer is known to attenuate the measurable SP signal (*36*). Nonetheless, this SP translates to an average ∼29.3% direct differences in biofilm EET between the N and S electrodes (see materials and methods). For context, genetic deletion of the entire *extHIJKL* operon, which encodes one of the primary outer membrane multiheme cytochrome conduits in *G. sulfurreducens*, results in a ∼13.7% reduction in electrode-respiration capacity due to the redundancy of EET conduits (*37*). Furthermore, for electrochemical context, the SP magnitude reported here is on par with the kinetic variations previously observed when biofilms are cultivated on different crystallographic orientations of gold electrodes (*38*). Thus, aligning the acceptor spin with the CISS-preferred orientation yields a physiological enhancement on par with important genetic and material modifications.

Another indicator of the physiological relevance of this effect is that N vs. S asymmetry appears prominently in cyclic voltammetry (CV) under active metabolism (turnover conditions; Figs. 2 and 3) but is absent without acetate as an electron donor (non-turnover regime; fig. S3). This distinction aligns with the current understanding of the CISS effect as a dynamic transport effect rather than an equilibrium phenomenon (*39*–*42*). Continuous oxidation of acetate by cells throughout the biofilm pushes a directional electron flux through the chiral cytochrome network, actively delivering a spin-polarized current to the electrode. The driving force for this transport is the redox potential difference between the acetate oxidation and the electrode potential. In the absence of this metabolic driving force, the measured current reflects the Faradaic charging and discharging of the biofilm’s finite redox cofactor pool by the electrode. Because interfacial electron transfer between the electrode and biofilm is fast relative to both the scan rate and electron hopping through the biofilm’s redox network (*43*), the electrode-biofilm interface is maintained in a state of quasi-equilibrium without an active electron supply from acetate oxidation. Under these conditions, measured current reflects the thermodynamic charging state of the biofilm rather than driven transport, and the CISS effect is masked. The intrinsically dynamic nature of the spin-dependent respiration phenomenon is further illustrated by our *in situ* modulation experiments, performed under turnover conditions but on the same biofilm as the magnetization of the electrode is reversed (Fig. 4 and fig. S4). Altering the spin state of the electrode instantaneously modulates the respiratory flux on a faster timescale than any cellular regulatory or growth response could occur.

These findings provide the first *in vivo* demonstration that the chirality induced spin selectivity effect actively gates biological electron transfer. While spin-selective kinetics have been demonstrated in abiotic catalysis (*44*), their physiological role during active cellular metabolism had remained experimentally inaccessible. By leveraging the extracellular respiration of electrodes as a direct probe, we establish electron spin as a fundamental physical parameter of biological energy conservation. Ultimately, these results reveal a previously unrecognized quantum feature of cellular electron flux, providing a strategy to exploit spin-selective pathways for bioelectronics.

## Supporting information

Supplementary Materials for Spin-Dependent Extracellular Respiration

## Acknowledgments

We thank Ron Naaman for insightful conversations about the CISS effect. We also acknowledge the Translational Imaging Center at the University of Southern California for their support and the use of their instrumentation, and Katya Kadyshevskaya for assistance with graphics.

