## Supplementary Materials for Spin-Dependent Extracellular Respiration for "Spin-Dependent Extracellular Respiration"

[1]Department of Physics and Astronomy, University of Southern California, Los Angeles, CA, USA
[2]Institute of Physics, Polish Academy of Sciences, Warsaw, Poland
[3]Department of Biological Sciences, University of Southern California, Los Angeles, CA, USA
[4] Department of Chemistry, University of Southern California, Los Angeles, CA, USA
[†]These authors contributed equally to this work
[‡]Present address: Omenn-Darling Bioengineering Institute, Princeton University, Princeton, NJ, USA
[§]Present address: Wells Fargo, Charlotte, NC, United States

### Materials and Methods

### Microbial culture and pre-growth procedures

*Geobacter sulfurreducens* PCA strain was used in this study. All procedures were conducted under aseptic and anoxic conditions at a controlled temperature of 30 °C, unless stated otherwise. A defined medium (referred to as NB) containing (per L of distilled water) 0.38 g KCl, 0.2 g $NH_4Cl$, 0.069 g $NaH_2PO_4$. $H_2O$, 0.04 g $CaCl_2$. $2H_2O$, 0.2 g $MgSO_4$. $4H_2O$, supplemented with 1.64 g (20 mM) sodium acetate (NBA), and 4.64 g (40 mM) sodium fumarate (NBFA), were used for this study. For strain pre-growth, NBFA was used; for metabolic electrochemical experiments, NBA was used; and for non-metabolic electrochemical experiments, NB was used.

To start strain culture, semisolid NBFA agar plates were prepared by adding 1% (v/v) agar to the NBFA. NBFA agar plates were streaked with *G. sulfurreducens* from frozen stock and incubated for 10-14 days until fully grown colonies (~1 mm diameter) were observed. Culture growth conditions and procedures were identical to those described previously (*45*). Briefly, single colonies were plucked and cultivated in 1 ml liquid NBFA in 1.5 ml microcentrifuge tubes for ~3 days. The 1ml liquid culture was then inoculated into anaerobic serum tubes (Tube 1) containing 10 ml NBFA. Upon reaching the stationary phase ($OD_{600nm}$ 0.6-0.7), 1 ml of culture from Tube 1 was used to inoculate a second set of anaerobic tubes (Tube 2) containing 10 ml NBFA. Once Tube 2 reached exponential phase ($OD_{600nm}$ 0.4-0.5), 10 ml of this culture was used to inoculate the electrochemical reactors. Reactor media was supplemented with 0.4 mL of 0.5 M acetate to ensure a starting concentration of 20 mM acetate in the reactor.

### Ferromagnetic electrodes

The ferromagnetic (FM) electrodes were prepared as described previously (*14*, *15*). Briefly, the epitaxial nanostructures were grown using a molecular beam epitaxy (MBE PREVAC) system. The layers deposited on a Sapphire $Al_2O_3$ (0001) substrate consisted of Pt 5 nm/Au 20 nm/Co 1.5 nm/Au 5 nm, which were used as FM working electrodes for electrochemical experiments. These films present perpendicular anisotropy with a coercive field of 117 G. The rectangular-shaped hysteresis loop, measured using a Polar magneto-optic Kerr effect (P-MOKE) measurement, confirms the unidirectional alignment of spins in the cobalt film, perpendicular to the surface, indicating a stable magnetization up to the coercive field, beyond which a magnetization reversal occurs, switching between spin up (N) and spin down (S). The top gold (Au 5 nm) layer acts as a capping layer to prevent oxidation of the Co layer while providing a biocompatible interface for biofilm attachment.

**Electrochemical reactors**

Single-chambered electrochemical reactors consisting of a glass cone (BASi MF-1084) sealed airtight with a rubber O-ring and a custom polyether ether ketone (PEEK) plastic lid were used. The lid includes separate ports for electrodes, gas exchange, and sampling. Reactors were pre-cleaned with an acid-base treatment followed by autoclaving, as described elsewhere (*45*). An Ag/AgCl (1 M KCl) reference electrode anchored through a $Na_2SO_4$ salt-bridge (0.1M in 1% agar) was used to monitor the electrode potential throughout the electrochemical experiments. A Pt wire (4-5 cm long, 0.25 mm diameter) served as the counter electrode. Two oppositely pre-magnetized (N and S) FM electrodes served as the working electrodes, whereas two similarly pre-magnetized (N–N) FM electrodes were used for the control.

In preparation for each experiment, the FM electrodes were treated sequentially with acetone, isopropanol, distilled $H_2O$, and dried with sterile high-pressure nitrogen gas. Electrodes were connected to wires using silver paint. Silver paint connections were subsequently covered with a non-conductive, electrochemically inert, and biocompatible epoxy glue (3M™ Scotchcast flame-retardant electrical insulating resin 2131) and cured for 24 hours. Before installation, the FM electrodes were sterilized by dipping into 10% bleach for 10 seconds and rinsed twice with 50 ml sterile DI water, followed by pre-magnetization in the desired orientation.

**Biofilm growth and Microbial Electrochemistry**

Electrochemical growth was employed to cultivate *G. sulfurreducens* biofilms on the FM electrodes (*45*). The FM electrodes were poised at fixed potentials (-0.225 or 0.175 V vs Ag/AgCl 1 M KCl), and the current produced by the biofilms was recorded using chronoamperometry (CA) at 10-second time intervals using a Squidstat potentiostat (Admiral Instruments, Inc.). The reactor contained NBA medium, lacking any electron acceptor apart from the FM electrodes, and with 20 mM acetate as the sole carbon source and electron donor. During growth experiments, the medium was stirred using a magnetic bead at ~1000 rpm to ensure uniform nutrient mixing and improve homogeneity. No measurable magnetic field was present at the location of the FM electrodes (as measured using a Hall device gaussmeter, PCE Americas Inc, PCE-MFM 3500 AC/DC Magnetic Field Meter device).

Cyclic voltammetry (CV) measurements were performed using a Squidstat potentiostat. CVs were recorded at three stages: (i) before inoculation in sterile anaerobic NBA medium without cells (abiotic condition); (ii) when the CA current reached a maximum current density (turnover condition); and (iii) after the media was replaced with NB (non-turnover) media and kept overnight in CA conditions to reach negligible catalytic current production in the absence of acetate. For all conditions, three consecutive CV cycles were collected over a potential range of –0.6 to +0.4 V vs Ag/AgCl at a 1 or 10 mV $s^{-1}$ scan rate, and the last CV cycle was used for the data analysis.

**Microscopy and biofilm thickness characterization**

After electrochemical experiments, biofilms grown on FM electrodes were fixed directly on the electrode surface inside the electrochemical reactor using 2% paraformaldehyde for 30 min. Post fixation, FM electrodes with attached biofilms were removed and washed with 1X PBS to remove residual fixative. The electrodes were imaged under a dissection microscope, and the biofilm covered surface area was quantified using a custom Python script that creates a grayscale image and thresholds the edge of the lighter areas to segment it. The pixel area in the segmented region was calculated and converted into surface area. This active area was used to normalize the produced currents and determine the current density.

Following imaging, biofilms were stained with the fluorescent membrane stain FM4-64FX (0.25 ug/ml) and incubated in the dark for 30 minutes. Excess stain was removed by gentle 1X PBS washing, and biofilms were imaged with confocal laser scanning microscopy (Zeiss LSM 780) integrated with an AxioObserver.Z1 inverted microscope (Carl Zeiss, GmbH, Jena, Germany). Images were captured using a 40× objective with an excitation at 561 nm and emission collected in a red channel (~600–750 nm) to capture the membrane fluorescence signal. Z-stacks were collected in at least three representative regions per electrode across the entire biofilm thickness with an appropriate z-step to ensure accurate 3D reconstruction.

***In Situ* Modulation of Respiratory Current Experiments**

For *in situ* magnetization reversal, the electrochemical reactors and pre-growth conditions were identical to those described in previous sections. To achieve a fully developed biofilm, the FM electrodes were poised at a constant applied potential (0.35 V vs. Ag/AgCl 1M KCl). Once the CA, $I(t)$, reached maximum current density, the magnetization of the FM electrode was reversed every 20 minutes (S to N or N to S) by momentarily bringing an external neodymium magnet (~1000 G surface magnetic field) close to the FM electrode and immediately removing it. The acquired signal, $I(t)$, was sequentially analyzed to quantify the modulation in EET flux following each magnetization reversal.

**Data analysis**

**Microbial Electrochemistry Direct Differences analysis**

The direct differences in biofilm EET between the N and S electrodes were calculated by subtracting $I_S$ from $I_N$ and normalizing by $I_S$, then multiplying by 100 to get a percentage value. The direct difference from reactors 1, 2, and 4 was then averaged, and the error is the standard deviation.

***In Situ* Modulation of Respiratory Current Experiments Analysis**

The raw current traces (fig. S5A) exhibit two superimposed components: i) a gradual monotonic increase in current over time due to progressive biofilm growth and increased bacterial colonization of the electrode surface, and (ii) periodic current modulations associated with controlled reversals of substrate magnetization as mentioned above.

To isolate the relevant component (ii), the long-timescale growth trend was removed by linear detrending. For each dataset, the current *I(t)* was fit to a first-order polynomial shown in eq. S1:

$$(eq.\ \ S1)\quad I_{fit}(t) = \ at + b$$

where $t$ represents elapsed time and $a$ and $b$ are the fitted slope and intercept, respectively. The detrended current was then computed as eq. S2:

$$(eq.\ \ S2)\quad I_{detrended}(t) = I(t) - I_{fit}(t)$$

This procedure removes the slow biofilm-driven drift while preserving the higher-frequency, magnetization-dependent current variations. Magnetization states were defined using time intervals corresponding to externally controlled magnetization orientation (N or S), which were identified from experimental timing records and applied consistently to both FM and control gold (Au) electrodes. For each magnetization interval, the detrended current was averaged to obtain a block-mean value. The change in current associated with each magnetization reversal was then calculated as the difference between the mean detrended current of adjacent blocks. Each current step was classified according to the magnetization reorientation direction (i.e., N to S or S to N), and summarized using the average of the step size and the error bars represent the full spread of the data. Steps were optionally excluded from analysis if predefined criteria were met (e.g., anomalous intervals), and exclusion criteria were applied independently to FM and Au datasets. Results were visualized as grouped bar plots comparing S to N and N to S transitions for FM and Au electrodes. Identical magnetization protocols were applied to both electrodes to ensure direct comparability. All data processing and analysis were performed in MATLAB (MathWorks), using custom scripts that extracted magnetization intervals, performed linear detrending, computed block averages, classified flip directions, and generated figures.

**Logistic growth model**

Current production was modeled using a logistic growth function (*29*),

$$(eq.\ \ S3)\qquad I(t) = \frac{I_{max}}{1 + \left(\frac{I_{max} - I_0}{I_{max}}\right) \cdot exp\,[-r \cdot t]}$$

where $I_{max}$ is the steady-state current density, $I_0$ is the initial current, and $r$ is the effective growth rate constant. Model parameters were obtained by nonlinear least-squares fitting using a MATLAB function (lsqcurvefit). This formulation assumes that current is proportional to electroactive biomass and that growth is limited by a carrying capacity corresponding to the maximum biofilm-supported current. After applying the model to the N and S magnetized data sets, temporal spin polarization (*SP(t)*) was calculated using eq. 2. Both measured *SP(t)* (from interpolated data) and predicted *SP(t)* (from fitted logistic models) were computed for reactors 1-4. Values were excluded when the denominator approached zero to avoid numerical instability.

To quantify growth dynamics, the numerical derivative was calculated from experimental data by interpolating currents onto a common time grid, applying a moving-average smoothing filter, and computing the derivative using a central-difference gradient operator. This provides an experimental estimate of instantaneous current change and by extension, the change in biofilm electron transfer resulting from an increase in biomass (*46*). Since this derivative approach is inherently noisy, the time derivative for the fitted current curves was also calculated analytically from eq. S4 as a function of $I(t)$:

$$(eq.\ \ S4)\ \ \frac{dI}{dt} = r \cdot I(t) \cdot (1 - \frac{I(t)}{I_{max}}\ )$$

**Nernst-Monod Model of Cyclic Voltammetry**

The CV measurements were modeled using a Nernst-Monod expression (*26*) with an added linear term to account for continued biofilm growth over time:

$$(eq.\ \ S5)\qquad I(V) = \frac{A}{1 + ex[-k \cdot (V - B)]} + C \cdot V$$

where $A$ represents the amplitude of the sigmoidal current response, $B$ is the midpoint (inflection potential), $C$ is a linear background slope reflecting continued biofilm growth during the CV measurements, and $k$ is a fixed steepness parameter $\frac{nF}{RT}$ (38.92 $V^{-1}$).

To find the experimentally relevant parameters for each CV measurement, the forward scan (increasing potential) was isolated by identifying the indices corresponding to the global minimum and maximum voltage values. The segment between these points was extracted and sorted to ensure monotonic increase in voltage. This procedure ensured consistent comparison of electrochemical behavior during the forward scan. The forward CV response was fitted to the Nernst-Monod model to extract $A, B, C$ via nonlinear least-squares fitting using a MATLAB function (lsqcurvefit). Potential-dependent spin polarization (*SP(E)*) was computed as a function of applied potential using eq. 1. To enable direct comparison, both datasets were interpolated onto a common voltage grid using piecewise cubic Hermite interpolation (pchip). Values were excluded when the denominator approached zero to avoid numerical artifacts. Spin polarization was calculated both from interpolated experimental data (measured *SP(E)*) and fitted model currents (predicted *SP(E)*), allowing assessment of model accuracy in capturing spin-dependent behavior.

**Biofilm Thickness Analysis**

Biofilm thickness was quantified using BiofilmQ (*47*). Confocal microscopy Z-stacks were segmented using automated global thresholding (RobustBackground method), with an identical thresholding sensitivity of 0.6 across all samples within each experiment. 3D reconstructions were generated based on voxel connectivity per 20 pixels, which correlates to 4.56 μm, and biofilm thickness was calculated as the vertical distance from the electrode surface to the outermost segmented biomass.

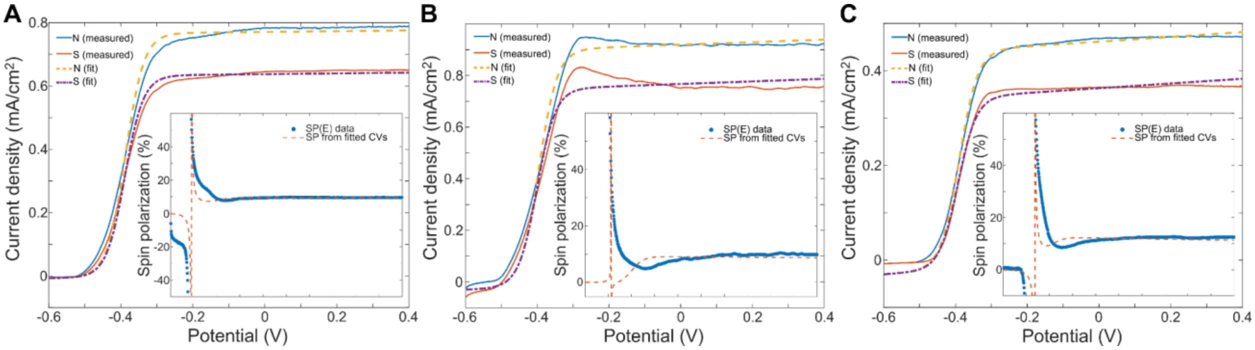


**Fig. S1: Nernst-Monod modelling of turnover cyclic voltammograms.** Nernst-Monod sigmoidal model fits (dashed lines) applied to the experimentally measured forward-scan CV data (solid lines) for Reactor 1 **(A)**, Reactor 2 **(B)**, and Reactor 3 **(C)**. (Insets) Expected potential-dependent spin polarization, *SP(E),* calculated from the analytical Nernst-Monod fits (dashed line) plotted against the experimentally measured *SP(E)* data (markers), the inset x-axis scale is identical to the main scale. The parameters extract distinct maximum limiting current amplitudes and minor shifts in midpoint potentials to capture the bioelectrochemical divergence between the N and S states.

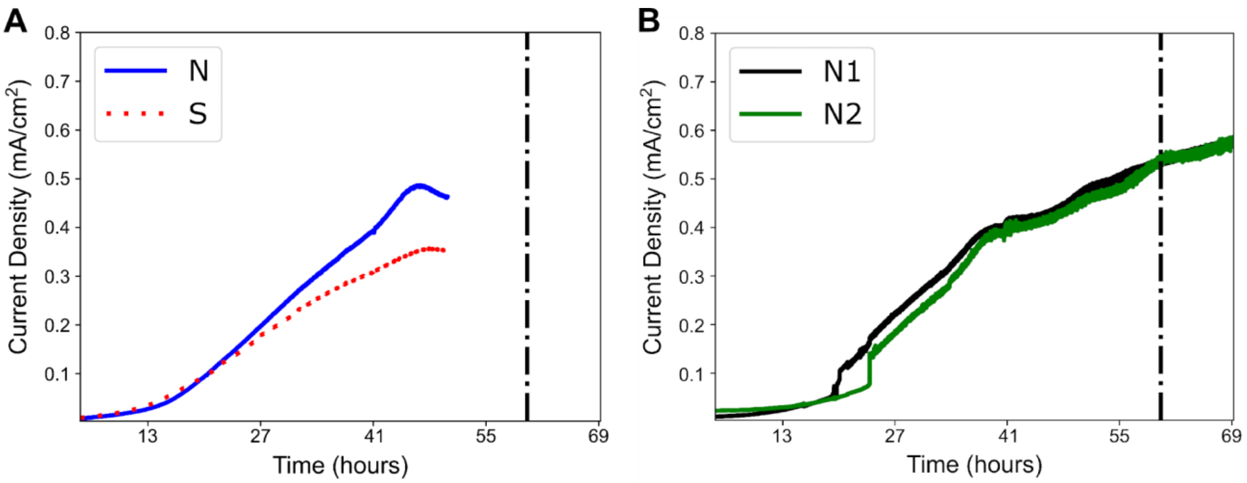


**Fig. S2: Chronoamperometric current profiles for partial-growth and identical-magnetization control reactors. (A)** Chronoamperometry, *I(t)*, of *G. sulfurreducens* biofilm growth on paired N and S electrodes in Reactor 3. The vertical dot-dashed black line represents the time at which the reactor was intentionally terminated prior to reaching maximum current density to allow for mid-growth biofilm analysis. **(B)** Chronoamperometry, *I(t)*, of a control reactor containing two identically magnetized (N-N) FM electrodes. Both electrodes display highly similar growth trajectories and steady-state currents, confirming that baseline reactor conditions do not inherently favor one electrode over the other in the absence of opposite magnetic orientation.

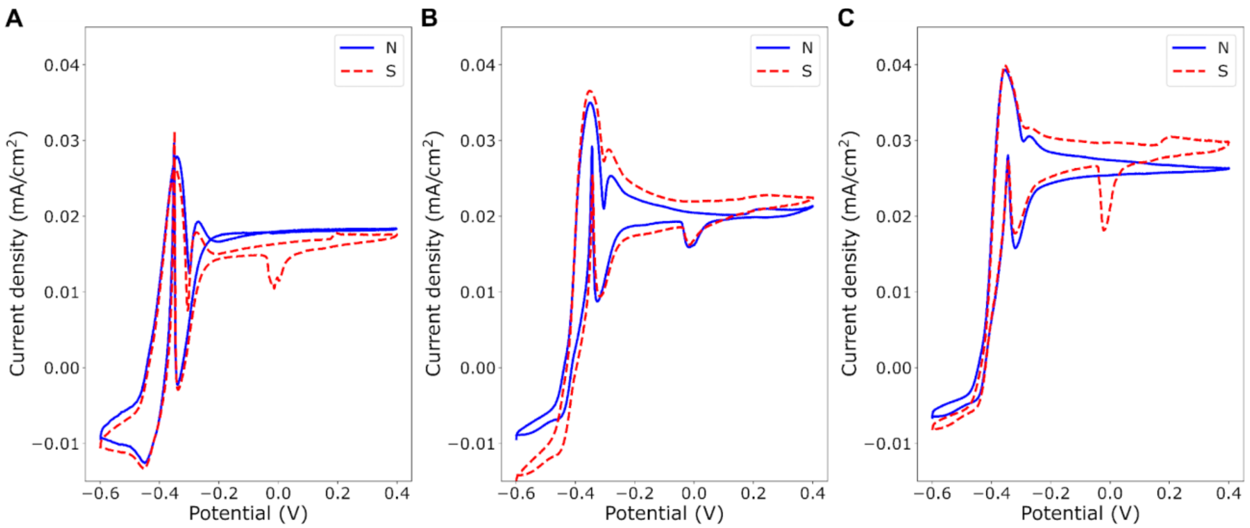


**Fig. S3: Non-turnover cyclic voltammetry of mature biofilms.** Non-turnover cyclic voltammograms (CVs) conducted at a slow scan rate of 1 mV/s following the replacement of standard growth media with acetate-free media. Panels **(A)**, **(B)**, and **(C)** display the CVs for Reactors 1, 2, and 3, respectively.

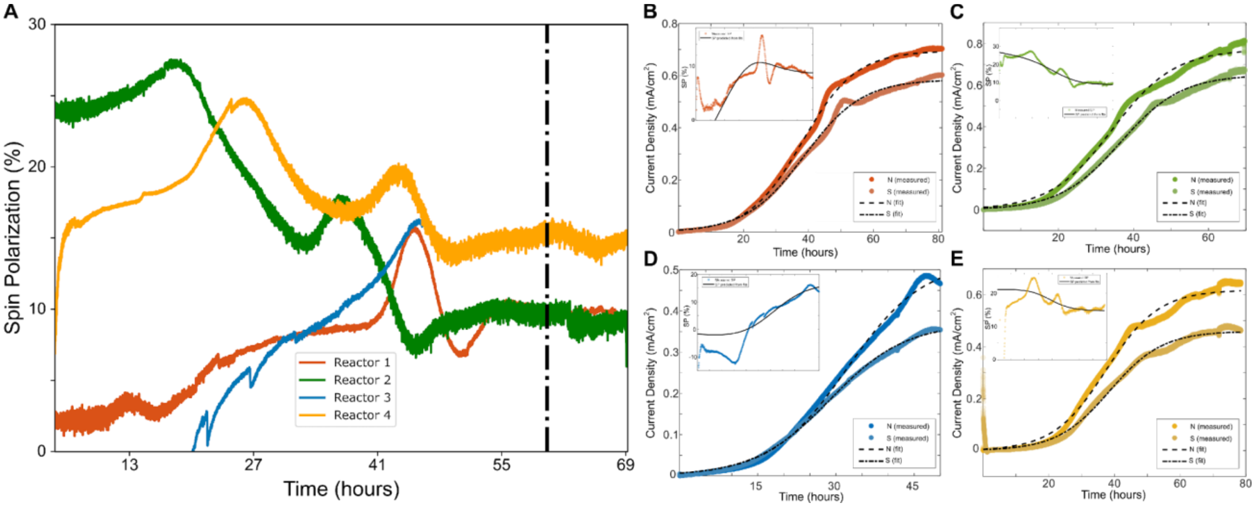


**Fig. S4: Temporal spin polarization and logistic growth modelling. (A)** Calculated temporal spin polarization, $\boldsymbol{SP}(\boldsymbol{t})$, derived from the raw chronoamperometric growth data across four reactors. **(B to E)** Logistic growth model fits (black dashed lines) overlaid on the experimental chronoamperometry data (colored markers) for Reactor 1 **(B)**, Reactor 2 **(C)**, Reactor 3 **(D)**, and Reactor 4 **(E)**. (Insets) *SP(t)* calculated from the analytical model fits (black line) plotted alongside the experimentally measured *SP(t)* (colored lines) for each reactor, the inset x-axis scale is identical to the main scale. Fitting parameters used to create the model fits are shown in table S4.

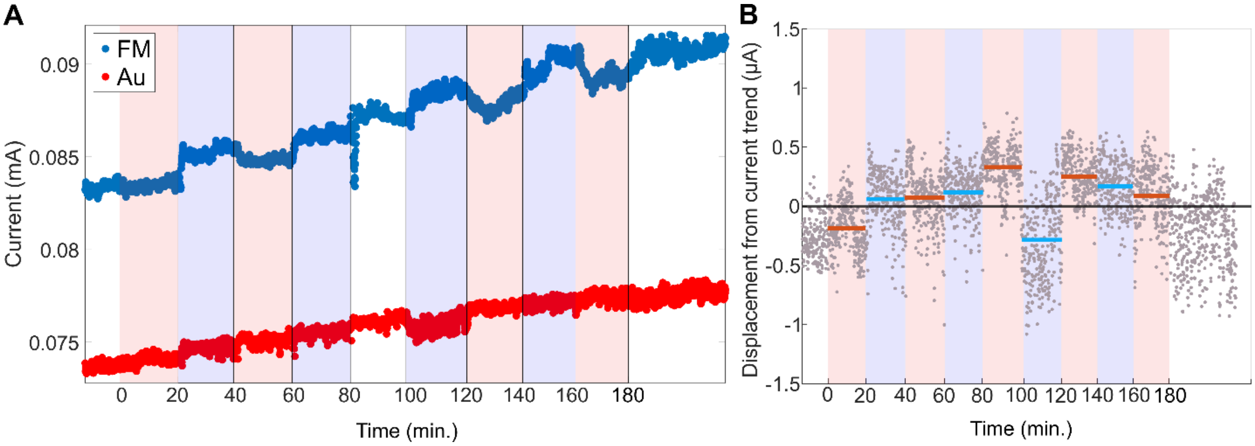


**Fig. S5: Raw in situ modulation data and non-magnetic control.** **(A)** Raw in situ modulation of respiratory current from biofilms on FM electrode (blue) and a non-magnetic gold (Au) control electrode (red). The background shading indicates the time intervals where the applied remanent magnetization state is N (blue shading) or S (red shading). The white background interval (**8**0 to **10**0 minutes) denotes an excluded period **in the FM electrode** due to a mechanical disturbance to the reactors that affected the current production at that time frame. **(B)** Detrended current profile for the Au control experiment. The linear biological growth trend was removed to isolate variations corresponding to magnetization reversal events. Horizontal bars indicate the average current for each 20-minute interval.

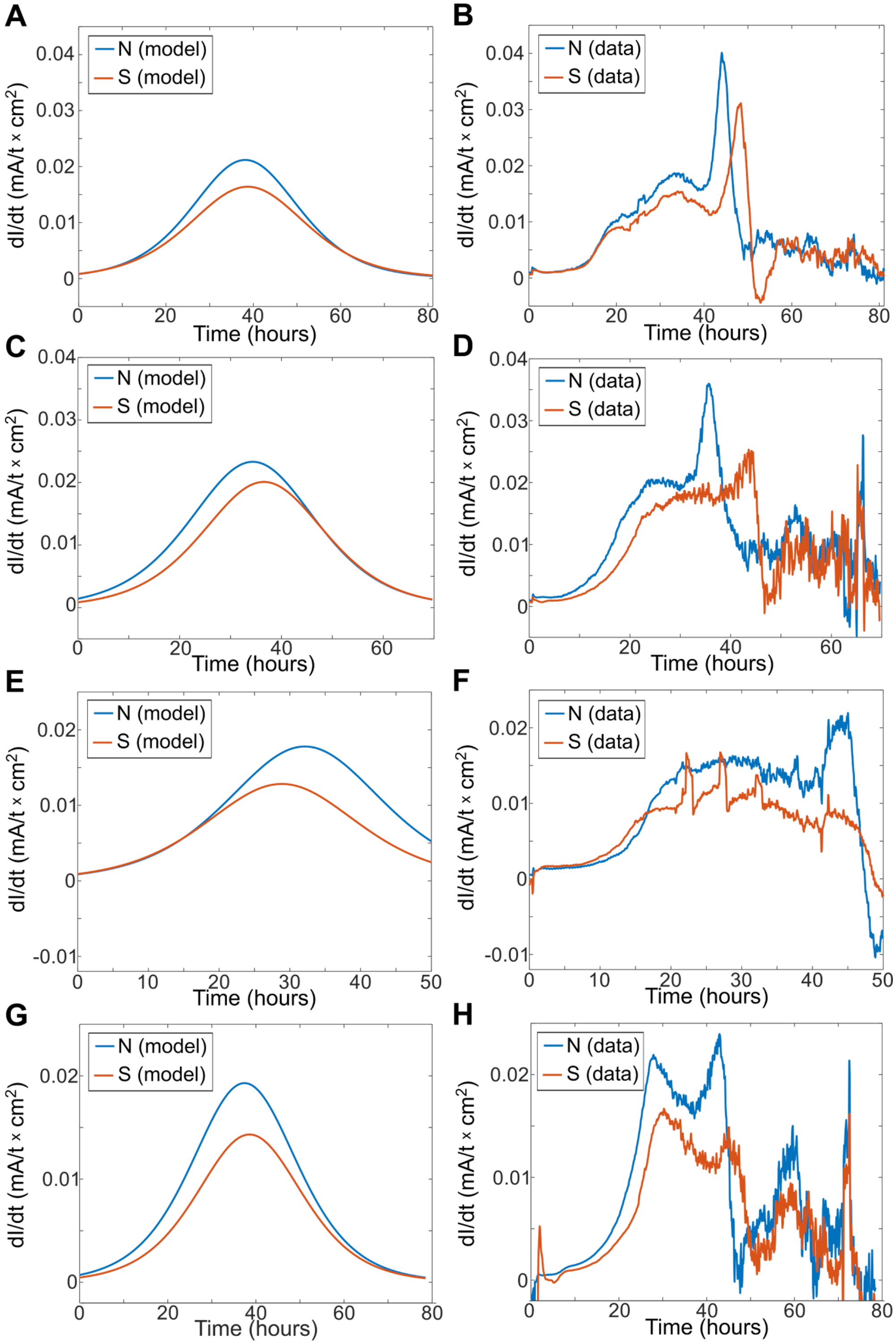


**Fig. S6: Time derivative of biofilm currents.** Each row **(A** and **B**, **C** and **D**, **E** and **F**, and **G** and **H)** corresponds to an independent biological reactor (Reactors 1 to 4, respectively). The left column **(A**, **C**, **E**, and **G)** displays the analytical time derivative calculated from the logistic growth model fits. The right column **(B**, **D**, **F**, and **H)** displays the numerical time derivative computed directly from the smoothed experimental chronoamperometric data. Blue lines denote the N-magnetized electrodes; red lines denote the S-magnetized electrodes.

**Table S1: List of replicate reactors and their relevant parameters.**

| **Reactor Number** | **Applied Potential for Growth** (V vs. Ag/AgCl 1M KCl) | **Rate of CV at the End of Growth** $\left(\frac{mV}{sec}\right)$ | **Notes** |
|---|---|---|---|
| Reactor 1 | 0.175 | 1 | |
| Reactor 2 | 0.175 | 10 | |
| Reactor 3 | 0.175 | 1 | Stopped before reaching a maximum current density |
| Reactor 4 | -0.22 | - | A faulty reference electrode led to a different potential during growth. |
| Control N-N | 0.175 | 10 | |

**Table S2: Fitting parameters for the Nernst-Monod Model (eq. S5) of Cyclic Voltammetry.** Where $\boldsymbol{A}$ represents the amplitude of the sigmoidal current response, $\boldsymbol{B}$ is the midpoint (inflection potential), $\boldsymbol{C}$ is a linear background slope reflecting continued biofilm growth during the CV measurements, and $\boldsymbol{R^2}$ is the statistical coefficient of determination. The forward CV response was fitted to the Nernst-Monod model via nonlinear least-squares fitting.

| **Reactor** | **Magnetization** | $\boldsymbol{A\left(\frac{mA}{cm}\right)}$ | $\boldsymbol{\Delta A}$- $\boldsymbol{(A_N - A_S)}$ | $\boldsymbol{B\ (mV)}$ | $\boldsymbol{\Delta B}$- $\boldsymbol{(B_N - B_S)}$ | $\boldsymbol{C\left(\frac{mA}{cm \cdot mV}\right)}$ | $\boldsymbol{R^2}$ |
|---|---|---|---|---|---|---|---|
| Reactor 1 | North | 0.77 | 0.13 | -388.5 | 1.6 | 0.027 | 0.9945 |
| | South | 0.64 | | -390.1 | | 0.025 | 0.9969 |
| Reactor 2 | North | 0.92 | 0.15 | -395.4 | 5 | 0.050 | 0.9917 |
| | South | 0.77 | | -400.4 | | 0.050 | 0.9864 |
| Reactor 3 | North | 0.46 | 0.1 | -389.4 | 3.2 | 0.050 | 0.9972 |
| | South | 0.36 | | -392.6 | | 0.050 | 0.9950 |

**Table S3: Confocal microscopy and average thickness of *G. sulfurreducens* biofilms.** Representative two-dimensional thickness maps extracted from confocal z-stacks for biofilms cultivated on N-magnetized and S-magnetized electrodes in the same reactors. Bar plots quantifying the average biofilm thickness for the N (blue) and S (red) electrodes in each respective reactor. Error bars represent the standard deviation of the average thickness over all measured areas of each sample.

<table>
<tr><th></th><th>Reactor 1</th><th>Reactor 2</th><th>Reactor 3</th><th>Reactor 4</th></tr>
<tr><td rowspan="4">N</td>
<td>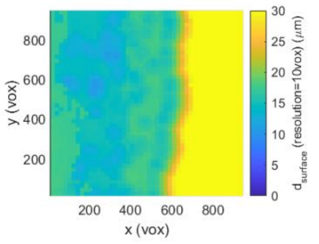</td>
<td>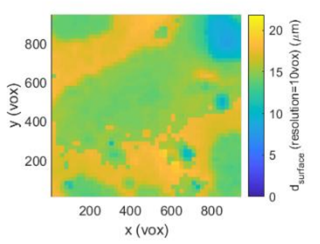</td>
<td>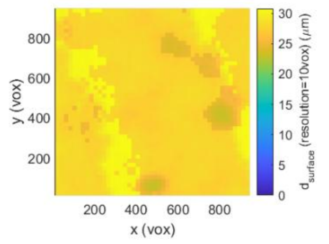</td>
<td>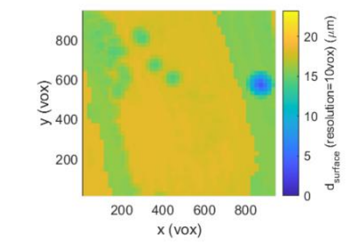</td></tr>
<tr>
<td>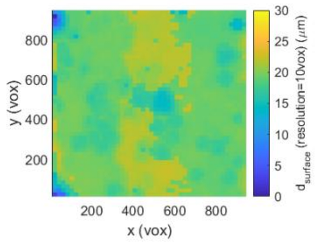</td>
<td>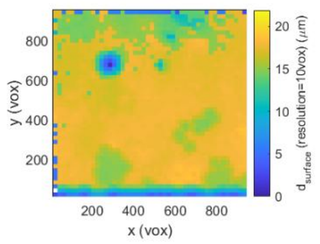</td>
<td>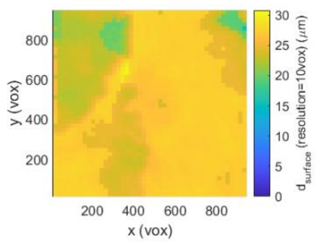</td>
<td>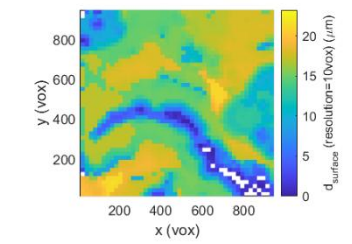</td></tr>
<tr>
<td rowspan="2">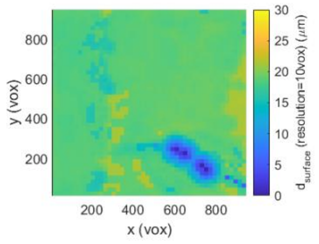</td>
<td>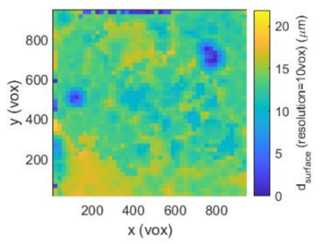</td>
<td rowspan="2">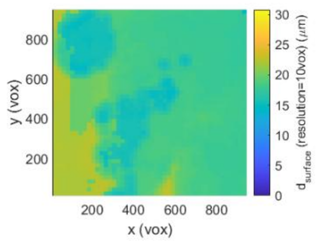</td>
<td rowspan="2">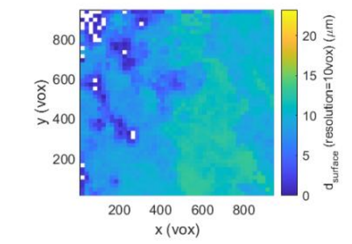</td></tr>
<tr>
<td>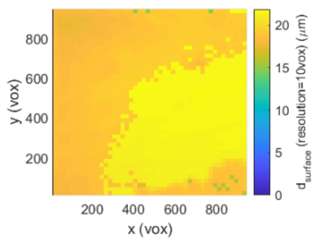</td></tr>
</table>

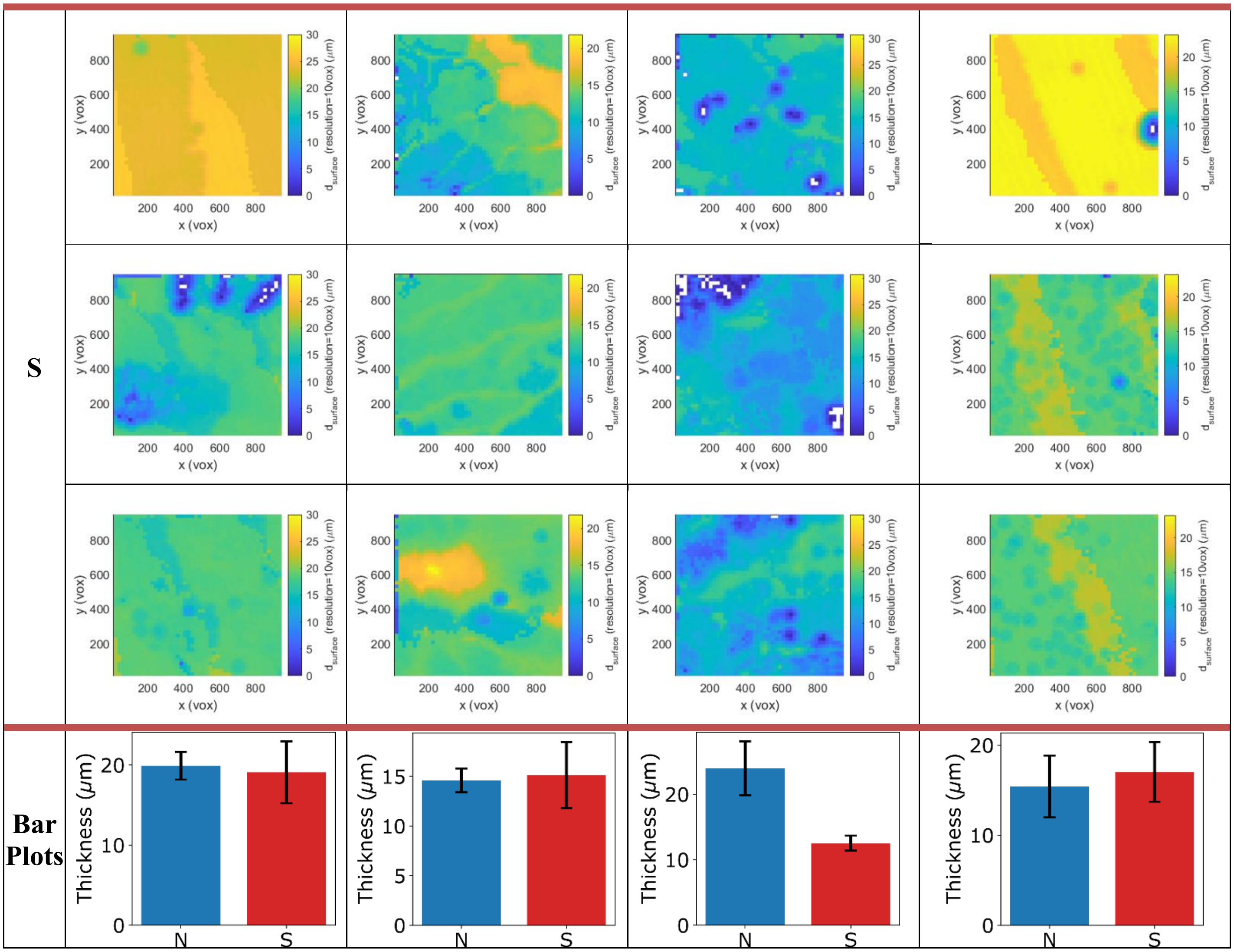
S
Bar Plots
x (vox)
y (vox)
dsurface (resolution=10vox) (μm)
Thickness (μm)
N
S

**Table S4: Fitting parameters for the logistic growth model (eq. S3).** Where $I_{max}$ is the steady-state current density, $I_0$ is the initial current, $r$ is the effective growth rate constant, and $R^2$ is the statistical coefficient of determination. Model parameters were obtained by nonlinear least-squares fitting.

| Reactor | Magnetization | $I_{max}\left(\frac{mA}{cm}\right)$ | $\Delta I_{max}$- $I_{max}^N - I_{max}^S$ | $I_0\left(\frac{mA}{cm}\right)$ | $\Delta I_0$- $I_0^N - I_0^S$ | $r\left(\frac{1}{hours}\right)$ | $R^2$ |
|---|---|---|---|---|---|---|---|
| Reactor 1 | North | 0.695 | 0.11 | 0.0066 | -0.0009 | 3.387e-5 | 0.9982 |
| | South | 0.585 | | 0.0075 | | 3.113e-5 | 0.9967 |
| Reactor 2 | North | 0.772 | 0.123 | 0.0120 | 0.0051 | 3.345e-5 | 0.9946 |
| | South | 0.649 | | 0.0069 | | 3.440e-5 | 0.9966 |
| Reactor 3 | North | 0.521 | 0.151 | 0.0064 | -0.0002 | 3.794e-5 | 0.9960 |
| | South | 0.370 | | 0.0066 | | 3.855e-5 | 0.9983 |
| Reactor 4 | North | 0.618 | 0.158 | 0.0057 | 0.0020 | 3.469e-5 | 0.9929 |
| | South | 0.460 | | 0.0037 | | 3.462e-5 | 0.9923 |
